# Sensorimotor dynamics differentiate singing and speaking

**DOI:** 10.64898/2025.12.19.695607

**Authors:** Alexis L. Pracar, Mattia F. Pagnotta, David R. Quiroga-Martinez, Rableen S. Ghuman, Cong Du, Mohammad Dastjerdi, Jack J. Lin, Jon T. Willie, Peter Brunner, Nina F. Dronkers, Robert T. Knight

## Abstract

Singing and speaking often dissociate clinically—people who stutter can sing fluently, and individuals with aphasia and speech output problems from stroke may express sentences fluently in song—yet the neural mechanisms of this centuries-old clinical phenomenon remain unclear. We recorded intracranial EEG while neurosurgical patients produced matched sentences by singing or speaking, sampling millimeter- and millisecond-scale activity across both hemispheres. During articulation, high-frequency activity (70–150 Hz) lateralized oppositely across behaviors, with right-dominant sensorimotor cortex (SMC) activation for singing and left-dominant activation for speaking. Phase-amplitude coupling revealed that mu-band (∼10 Hz) phase locally organized HFA in left SMC in a channel-specific, spatially interdigitated architecture preserved across both behaviors. Mu-band synchrony in speaking showed an early left-led pattern, whereas singing exhibited a ramping of synchrony within the left sensorimotor cortex and between the two motor cortices, supporting progressive interhemispheric recruitment. Frequency-domain Granger– Geweke causality revealed that the left primary somatosensory cortex drives both motor cortices at speech onset. In contrast in singing, control over motor cortices relied on both hemispheres. Singing and speaking engage a shared mu-coupled sensorimotor substrate through distinct recruitment dynamics.

## Introduction

For centuries, a striking dissociation has been noted between singing and speaking in many clinical populations. People who stutter when they speak do not stutter when they sing^1–3^. Individuals with aphasia, a language disorder occurring after brain injury, struggle to produce spoken utterances but fluently produce them through song^4–6^. Even in cases of dementia, singing familiar songs can induce moments of lucidity and boost cognitive function^7–9^. These observations have motivated the theory that speech and song production draw on distinct neural mechanisms, yet these mechanisms remain undefined. Given its central role in generating and monitoring articulatory gestures^10–16^, sensorimotor activity during singing provides a window into how singing diverges from speaking identical utterances.

We compared singing and speaking production in neurosurgical patients undergoing invasive monitoring for treating drug-resistant epilepsy, a clinical population that allows direct observation of real-time neural activity using intracranial electroencephalography (iEEG). Participants produced matched sentences in both modalities (singing and speaking) while we recorded millimeter-scale, millisecond-resolution activity across bilateral sensorimotor cortex (SMC) (Fig. 1). We quantified high-frequency activity (70–150 Hz; HFA), local phase-amplitude coupling (PAC), oscillatory synchrony (1–70 Hz), and directed connectivity during articulation to probe hemispheric and sensorimotor dynamics of speaking versus singing linking clinical observations to circuit-level physiology.

**Figure 1.**
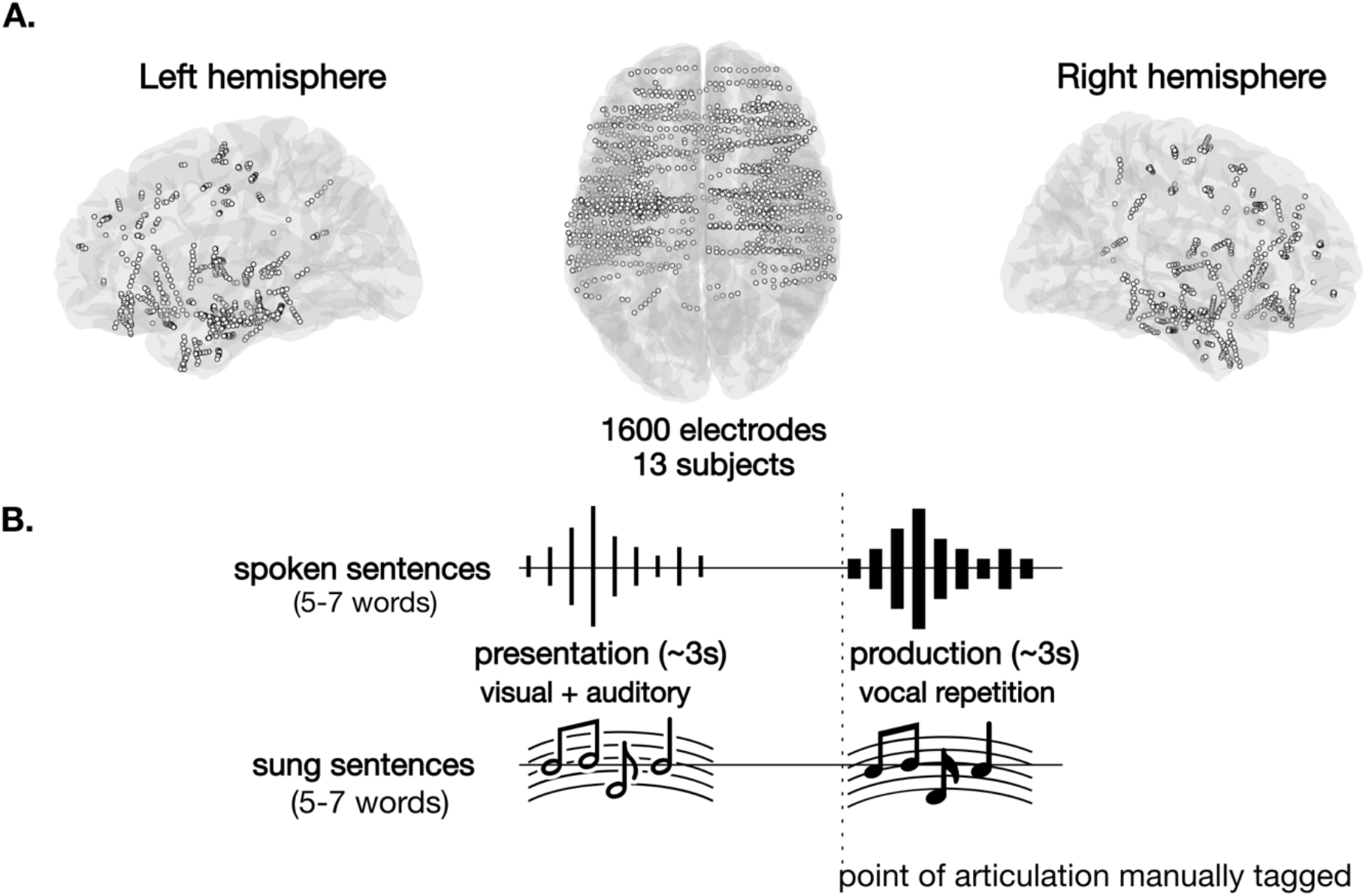
Task design and electrode coverage. **A**. Coverage of all electrodes (N=1600) across the subjects included in the final analysis (N=13) is shown in MNI standard space, including coverage of the left and the right hemisphere. **B**. Schematic representation of the task design. Matched sentences (20 per condition) were presented visually and auditorily to participants. Participants were then asked to repeat them once either while singing or while speaking (in alternate blocks), as quickly as possible. The stimuli were designed to engage motor speech planning and execution skills. The stimuli contained 5–7 words, some of which were multisyllabic with consonant clusters that required shifting between points of articulation (see Methods for details).

## Results

### High frequency activity and the mu rhythm in sensorimotor cortex

We observed higher articulation-locked HFA for singing in the right sensorimotor cortex (SMC; S1 and M1) and for speaking in the left SMC. Cluster-based permutations were run to directly compare singing versus speaking from 0.5 s before articulation onset throughout the first 2 seconds of production. Time-varying power estimates were derived using a Morlet wavelet approach during the production period following the onset of articulation, which was manually identified from the audio files for each recording session. High frequency activity (HFA; 70–150 Hz) is a key marker of local cortical activation^17,18^ and drives the BOLD signal^19,20^. In the left SMC (S1 and M1), HFA power was higher in the speaking condition (Left S1: *p*_*perm*_=0.0004; Left M1: *p*_*perm*_=0.0005) (Fig. 2A,B). In the right SMC (S1 and M1), HFA power was higher in the singing condition (Right S1: *p*_*perm*_=0.0006; Right M1: *p*_*perm*_=0.0002, 0.008) (Fig. 2C,D). Regions of interest analyzed were these structures bilaterally: inferior frontal gyrus (IFG), superior temporal gyrus (STG), postcentral gyrus (PostCG/S1), and precentral gyrus (PreCG/M1). Only significant results are reported.

**Figure 2.**
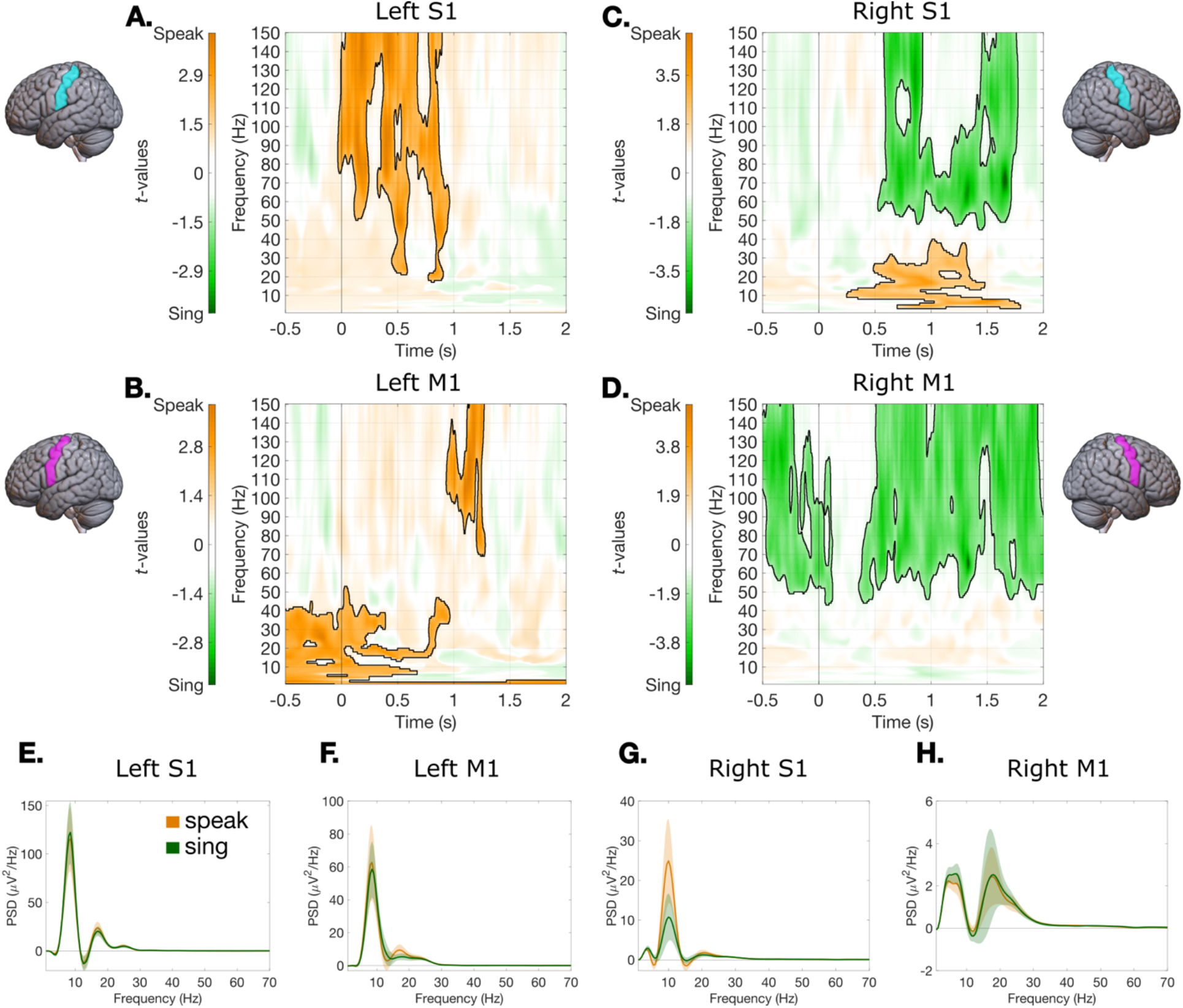
Time-varying power comparisons between speaking and singing after articulation. **A–D** Time-frequency distribution of the power differences (*t*-values) between speaking and singing. Cluster-based permutations were run over time and frequency bins (1–150 Hz; 10,000 permutations). Significant clusters with higher power for the speaking condition are shown in orange with a black outline. Significant clusters with higher power for the singing condition are shown in green with a black outline. **A&B**: In the left sensorimotor cortex (M1 and S1), higher HFA power is found in the speaking condition. **C&D:** In the right sensorimotor cortex (M1 and S1), higher HFA power is found in the singing condition. **E–H:** A mu rhythm (∼10 Hz) is visible in all areas, detected using Irregular-Resampling Auto-Spectral Analysis (IRASA),^29^ to separate fractal components (related to 1/f aperiodic activity) and oscillatory components in the power spectrum. In each panel, the shadings represent 95% confidence intervals (CIs) across the electrodes in that area.

Information transfer in a network is supported by low frequency oscillatory activity and coherence in the 1–70 Hz range^21–26^. Fluent output depends on millisecond-scale, precisely-timed motor commands that recruit dozens of muscles across the respiratory, laryngeal, and supralaryngeal systems to orchestrate coordinated tongue, lip, and jaw gestures that shape dynamic vocal-tract constrictions. In natural speech, transitions between these gestures are rapid and seamless^10,27,28^. Our power analysis also revealed lower-frequency rhythms in SMC (Fig. 2). Clusters in left and right SMC had higher power for the speaking condition in the 10–30 Hz range (*p*_*perm*_=0.0154, 0.0080) (Fig. 2B,C). We used Irregular-Resampling Auto-Spectral Analysis (IRASA)^29^, to separate fractal components (related to 1/f aperiodic activity) and oscillatory components in the power spectrum. A ∼10 Hz “mu rhythm” (with a 20 Hz harmonic) was prominent in SMC (Fig. 2E–H). The mu rhythm has been previously associated with speech and somatomotor activity^30– 32^. Together, the findings in Fig. 2 establish two features of sensorimotor activity during sentence production: HFA, which lateralizes oppositely for the two behaviors, and a prominent SMC mu rhythm, which is shared bilaterally across them.

### Mu phase coordinates high frequency activity in sensorimotor cortex

Having identified two co-existing signatures of SMC activity, a behavior-differentiating HFA response and a shared bilateral mu rhythm, we next asked whether they are mechanistically linked at the local level. HFA reveals which sensorimotor regions activate during articulation, but fluent vocal production also requires that this activation be precisely timed. To test whether sensorimotor HFA amplitude is organized in time by an underlying oscillatory rhythm, we quantified phase-amplitude coupling (PAC) between low-frequency phase and HFA amplitude, using Tort’s modulation index (MI)^33^ computed within the production window (0 to +2000 ms post-articulation) separately for each electrode and condition.

Across the group, mu-band phase organized HFA predominantly in the left sensorimotor cortex (Fig. 3A–C). When MI(z) was averaged across HFA frequencies and channels within each left-hemisphere ROI and plotted across phase frequencies, a clear peak emerged at ∼8–10 Hz in left S1 and left M1, but not in other canonical left language ROIs (STG, IFG) or adjacent ROIs (MFG, SMG) (Fig. 3A). The proportion of channels showing significant mu–HFA PAC was strongly left-SMC dominant: ∼39% of left S1 channels and ∼18% of left M1 channels showed significant coupling, while right-hemisphere SMC and bilateral language ROIs all remained below ∼7% (Fig. 3B). Significant PAC channels were anatomically concentrated in left SMC (Fig. 3C). This coupling was present in both speaking and singing, indicating that mu–HFA PAC is a shared property of left SMC during sentence production rather than a feature that differentiates the two behaviors. The significant channels were in middle and dorsal SMC, not in ventral SMC.

**Figure 3.**
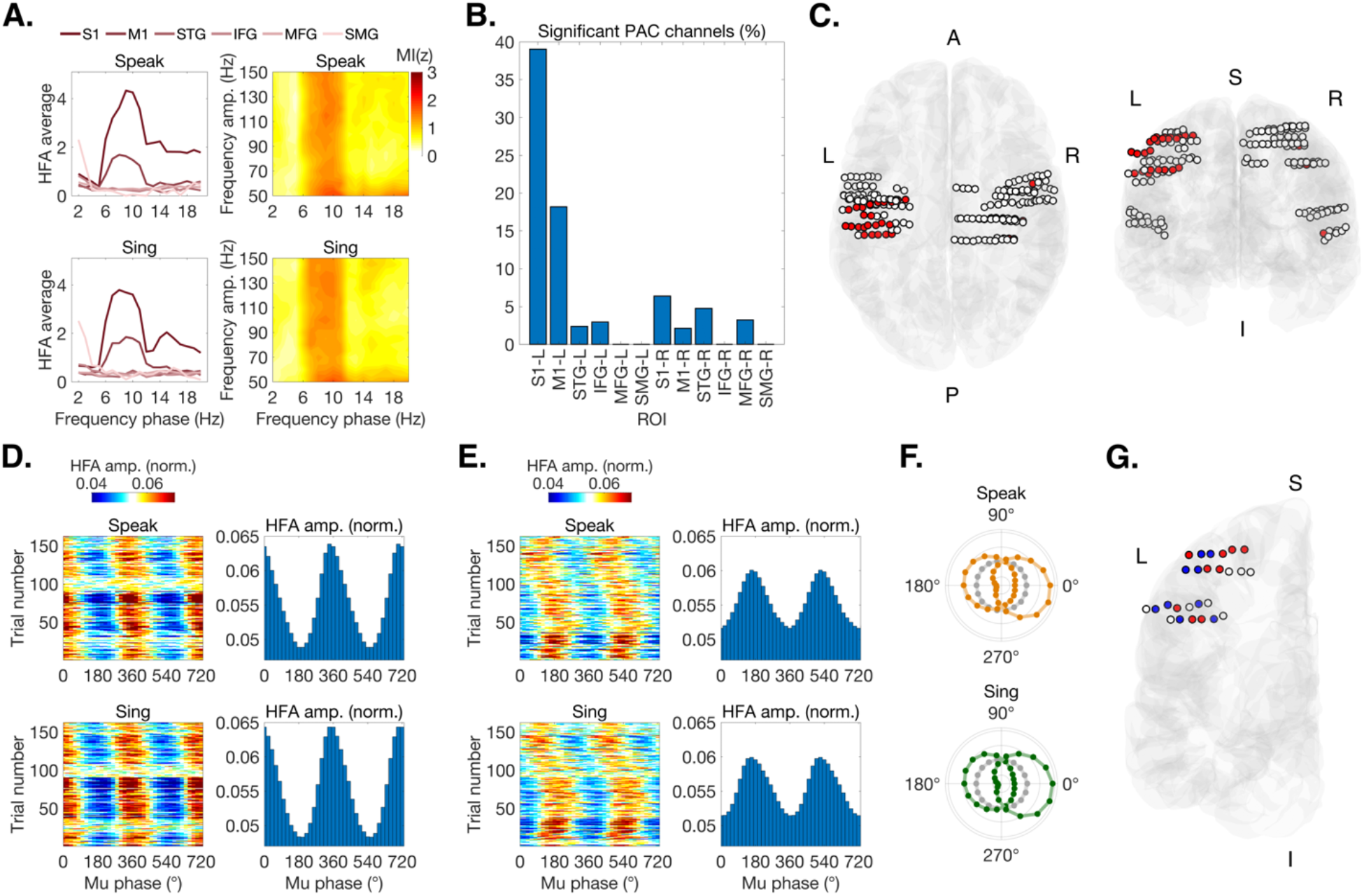
Mu-band phase organizes high-frequency activity in left sensorimotor cortex during sentence production. (**A**) Group-level z-scored MI (thresholded at z > 1.64, p < 0.05) averaged across HFA amplitude frequencies (70–150 Hz) and channels within each left-hemisphere ROI, plotted across phase frequencies (2–20 Hz), separately for speak (top) and sing (bottom). Both conditions show a clear MI peak at ∼8–10 Hz in S1 (dark red) and M1 (red), but not in language or adjacent ROIs (STG, IFG, MFG, SMG). Adjacent comodulograms (phase × amplitude frequency, MI(z)) confirm that coupling is concentrated on mu-phase × broadband HFA in both conditions. (**B**) Percentage of channels with significant mu–HFA PAC (mean MI(z) > 1.64 in the 8–10 Hz × 70–150 Hz window), per ROI × hemisphere, averaged across conditions. Left S1 (∼39%) and left M1 (∼18%) dominate; all other ROIs, including right SMC and bilateral language regions, remain below ∼7%. (**C**) MNI-template renderings (axial and coronal views) showing all sampled bilateral SMC channels (white), with significant mu–HFA PAC channels shown in red. Significant PAC channels concentrate in left SMC. (**D**) Single-subject example (S6). Left-SMC channels whose HFA amplitude peaks preferentially lock to mu phase near 0°/360°. Left: trial-by-trial HFA amplitude (normalized) across two cycles of mu phase (0–720°). Right: HFA amplitude as a function of mu phase, averaged across trials. Top = speak, bottom = sing. Phase preference is stable across trials and preserved across conditions. (**E**) Same layout as D, for left-SMC channels in S6 whose HFA amplitude peaks preferentially lock to mu phase near 180°. (**F**) Polar plots for S6 showing HFA amplitude binned across one mu cycle (18 phase bins). Colored markers and lines = left SMC (orange = speak, green = sing); grey = within-subject reference distribution from left language ROIs. Left SMC shows a bimodal distribution with peaks near 0° and 180° in both conditions; left language ROIs do not show this structure. (**G**) Sagittal MNI view (S6) of left-SMC channels colored by their preferred mu phase: red = ∼0°/360°, blue = ∼180°, white = other left-SMC channels. Channels with opposite phase preferences are spatially interdigitated across left SMC with no clear macroscopic segregation.

To examine the temporal and spatial precision of this coupling, we re-estimated MI on individual trials in an example participant (S6), with broad electrode coverage within the left SMC (S1: 14 electrodes; M1: 12 electrodes). Within left SMC, individual channels showed PAC with HFA that was stable across trials and preserved across speaking and singing (Fig. 3D–E). Across left-SMC channels in this participant, preferred mu phase formed a bimodal distribution with peaks near 0° and 180°, a structure that was absent in other regions in canonical left language cortex (Fig. 3F). Channels with opposite phase preferences (0° or 180°) were spatially interdigitated across left SMC with no clear macroscopic segregation (Fig. 3G). Together with the trial-by-trial stability shown in Fig. 3D–F, this pattern indicates that mu-phase preference is a precise, channel-specific dynamic property of left SMC rather than a coarse anatomical gradient, consistent with prior ECoG work showing functionally specific but spatially intermixed representations in sensorimotor speech cortex^10,12^.

### Different mu-synchrony patterns arise in sensorimotor cortex during singing and speaking

The PAC results showed that mu phase locally organizes HFA within left SMC, in a manner shared by speaking and singing. Fluent vocal production, however, also requires precise coordination across sensorimotor sites and between the two hemispheres. We therefore examined pre- and post-articulatory synchrony dynamics in the left and right hemisphere for singing versus speaking. Regions of interest analyzed included these structures bilaterally: IFG, STG, MFG, SMG, S1, and M1. Only significant results are reported. Low frequency oscillatory synchrony further distinguished singing versus speaking. Our results revealed that singing evokes mu rhythm synchrony intra-hemispherically within the left SMC and inter-hemispherically between left and right SMC. Synchrony of intra- and inter-hemispheric regions was measured using intersite phase clustering (ISPC^34^), a measure of phase-based connectivity that quantifies phase-locking of signals over time. For each connection between two areas of interest, only subjects with electrode contacts in both areas were tested for synchronization between areas. Analysis was restricted to the oscillatory range (1–70 Hz) and cluster-based permutations were run on the ISPC estimates, across all pairs of electrode contacts between areas, to compare synchrony in singing and speaking. To capture the dynamics of preparation and articulation, we quantified the relative synchrony estimates in the mu rhythm (8–12 Hz) during singing and speaking compared to their pre-articulatory baselines (z-scored using 0.5 s before production).

#### The left sensorimotor cortex achieves speech-like mu-synchrony through singing

Left SMC mu-synchrony (between left M1 and left S1) peaked at speech onset (*p*_*perm*_=0.0020) but continued to ramp up during singing (*p*_*perm*_=0.0002), as revealed by directly comparing between conditions using a cluster-based permutation approach (Fig. 4A, left panel). For speaking, we observed stronger synchronization in the mu rhythm right before articulation, with an increase at the point of articulation, as shown by z-scored relative synchrony changes (RSC) in the mu-band (8–12 Hz), obtained by normalizing within-condition estimates using a 0.5 s pre-articulation baseline (Fig. 4A, right panel). On the other hand, for singing, we observed weaker synchronization at the onset, measured by RSC in the mu rhythm, however synchronization ramped up throughout production. The synchrony estimates also revealed that, in the left SMC, the mu rhythm is detectable in the speaking condition during the pre-articulatory period. In contrast, it is not present in the singing condition during the pre-articulatory period, but synchrony emerges during production. These observations indicate that singing achieves a sustained speech-like activity profile in the left mu rhythm without the same signatures of preparatory activity seen in speaking. Right SMC mu-synchrony was more sustained throughout the speaking condition (*p*_*perm*_=0.0020) and did not show a strong ramping up of synchrony in either condition (Fig. 4B).

**Figure 4.**
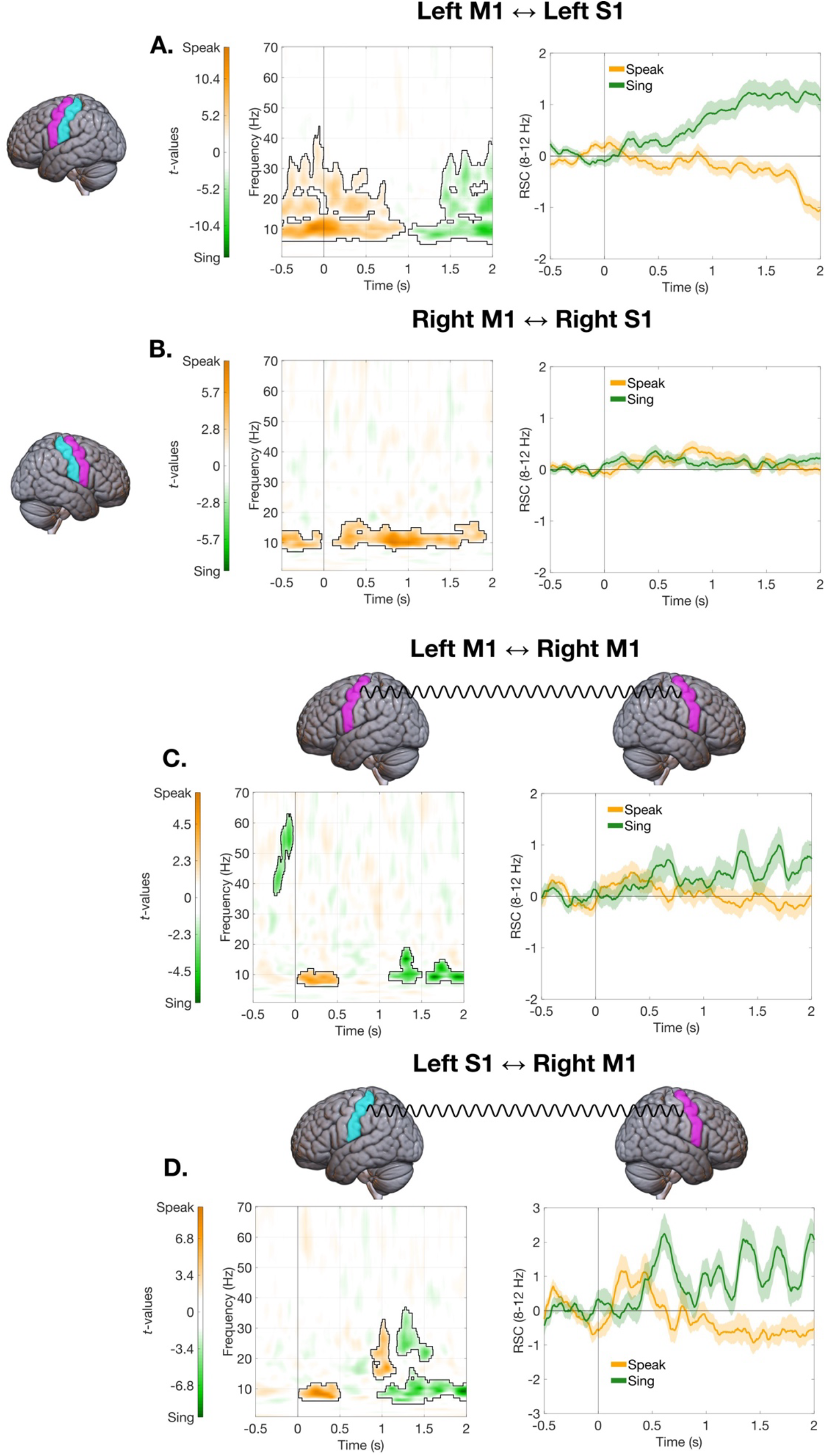
Synchrony ramps up in singing in the sensorimotor cortex. Time-frequency distribution of the intra- and interhemispheric synchrony differences (*t*-values) between speaking and singing. Synchrony estimates were derived using intersite phase clustering (ISPC). Cluster-based permutation was run on the pairs of contacts between areas in the 1–70 Hz range. Significant clusters with higher synchrony for the speaking condition are shown in orange with a black outline. Significant clusters with higher synchrony for the singing condition are shown in green with a black outline. Z-scored relative synchrony changes (RSC) in the mu-band (8–12 Hz), obtained by comparing the estimates in each condition to their own baselines of 0.5 s before articulation, are shown in the same color scheme (speak in orange, sing in green). **A:** In the left SMC (M1 ↔ S1), synchrony is initially higher in the speaking condition, but then is higher in singing as the sentence unfolds. **B:** In the right SMC (M1 ↔ S1), higher synchrony is found throughout the speaking condition. **C:** Left M1 ↔ Right M1. Between the left and right motor cortex (M1), initially synchrony is higher in the speaking condition, but synchrony ramps up during singing production. **D:** Left S1 ↔ Right M1. Between the left S1 and right M1, initially synchrony is higher in the speaking condition, but synchrony ramps up during singing production. In the right panels, the shadings represent 95% confidence intervals (CIs) across the electrode pairs for that connection.

#### Interhemispheric mu-synchrony increases in singing in sensorimotor cortex

Interhemispheric communication is integral to speech motor control. Left-hemisphere stroke with aphasia is often accompanied with right-sided facial drooping^35^, reflecting predominantly crossed control of facial and articulatory musculature^36,37^ (e.g., corticobulbar projections). In fluent production, the sensorimotor system must coordinate both hemispheres to drive symmetric, precisely-timed movements of lips, tongue, and jaw^10,12,27,38,39^. Differential singing modulation of left–right SMC interactions may explain how song can bypass or mitigate motor deficits after left-hemisphere injury.

Interhemispheric synchrony between the motor cortices (left M1–right M1) at the onset of articulation was initially higher for speaking (*p*_*perm*_=0.0119) (Fig. 4C) but ramped up during singing (*p*_*perm*_=0.0060, 0.008, 0.0125). Interhemispheric synchrony between left S1 and right M1 also showed higher mu synchrony at the onset of articulation for speaking (*p*_*perm*_=0.0004; 0.0009) (Fig. 4D). However, as the sentence unfolded, there was higher synchrony for singing (*p*_*perm*_=0.0002; 0.0006).

### Directional mu-band influences across sensorimotor cortex differ for singing and speaking

We observed distinct mu-band synchrony profiles in SMC for speaking and singing at articulation onset and later in production. Singing elicited overall higher mu-band synchrony as production unfolded, with different preparatory signatures. To probe network directionality, we applied Granger causality analyses to identify which SMC nodes lead during speaking vs. singing. Using a frequency-domain Granger–Geweke Causality (GGC) framework^40^, we estimated directed influences between areas in two windows (initiation: 0–0.5 s post-articulation onset and production: 1–1.5 s post-articulation onset). We first determined the dominant directionality of the influence between the two areas by testing if the net influence (area 1 → area 2 minus area 2 → area 1) differed significantly from zero, separately for each condition (speak and sing). This revealed if, regardless of condition, there was a net flow of information from one area to another, establishing a “controller” of the system. Then, in each inter-areal connection, we compared the mu-band (8–12 Hz) directed connectivity of speaking versus singing using a paired-samples *t*-test across all electrode pairs for that connection, to determine if connectivity was stronger in one condition.

#### Initiation: 0–0.5s post-articulation

The left sensory cortex S1 exerted influence on motor cortices in both hemispheres at initiation. In the left hemisphere, mu-band connectivity was higher for speaking in the connections from left S1 → left M1 (*t*(548)=9.6041, *p*<0.0001) and from left M1 → left S1 (*t*(548)=8.0833, *p*<0.0001) (Fig. 5A,C). The net influence direction was from left S1 → left M1 in both speaking (*t*(548)=-3.7870, *p*=0.0002) and singing (*t*(548)=-3.8665, *p*=0.0001) (Fig. 5C). Left S1 also led right M1 in both speaking (*t*(69)=5.4482, *p*<0.0001) and singing (*t*(69)=5.6322, *p*<0.0001) but mu-band connectivity between these two areas was higher in speaking compared to singing (left S1 → right M1: *t*(69)=7.2398, *p*<0.0001; right M1 → left S1: *t*(69)=6.1756, *p*<0.0001) (Fig. 5D). Between the two motor strips, mu-band connectivity was higher in speaking (left M1 → right M1: *t*(189)=5.4545, *p*<0.0001; right M1 → left M1: *t*(189)=4.8184, *p*<0.0001) (Fig. 5E). We found a stronger net influence direction from left M1 → right M1 in the speaking condition (*t*(189)=2.5321, *p*=0.0122), but not in the singing (*t*(189)=1.6623, *p*=0.0981). Even though we found that, in both speaking and singing, right S1 was leading right M1 (speaking: *t*(870)=-10.5858, *p*<0.0001; singing: *t*(870)=-8.4781, *p*<0.0001) and left M1 (speaking: *t*(142)=-4.3683, *p*<0.0001; singing: *t*(142)=-3.4998, *p*=0.0006) (Fig. 5F), we did not find any significant differences between conditions in net influence direction (right M1 → right S1: *t*(870)=-1.2078, *p*=0.2275; right S1 → right M1: *t*(870)=1.5970, *p*=0.1106; left M1 → right S1: *t*(142)=-0.9901, *p*=0.3238; right S1 → left M1: *t*(142)=0.1996, *p*=0.8420).

**Figure 5.**
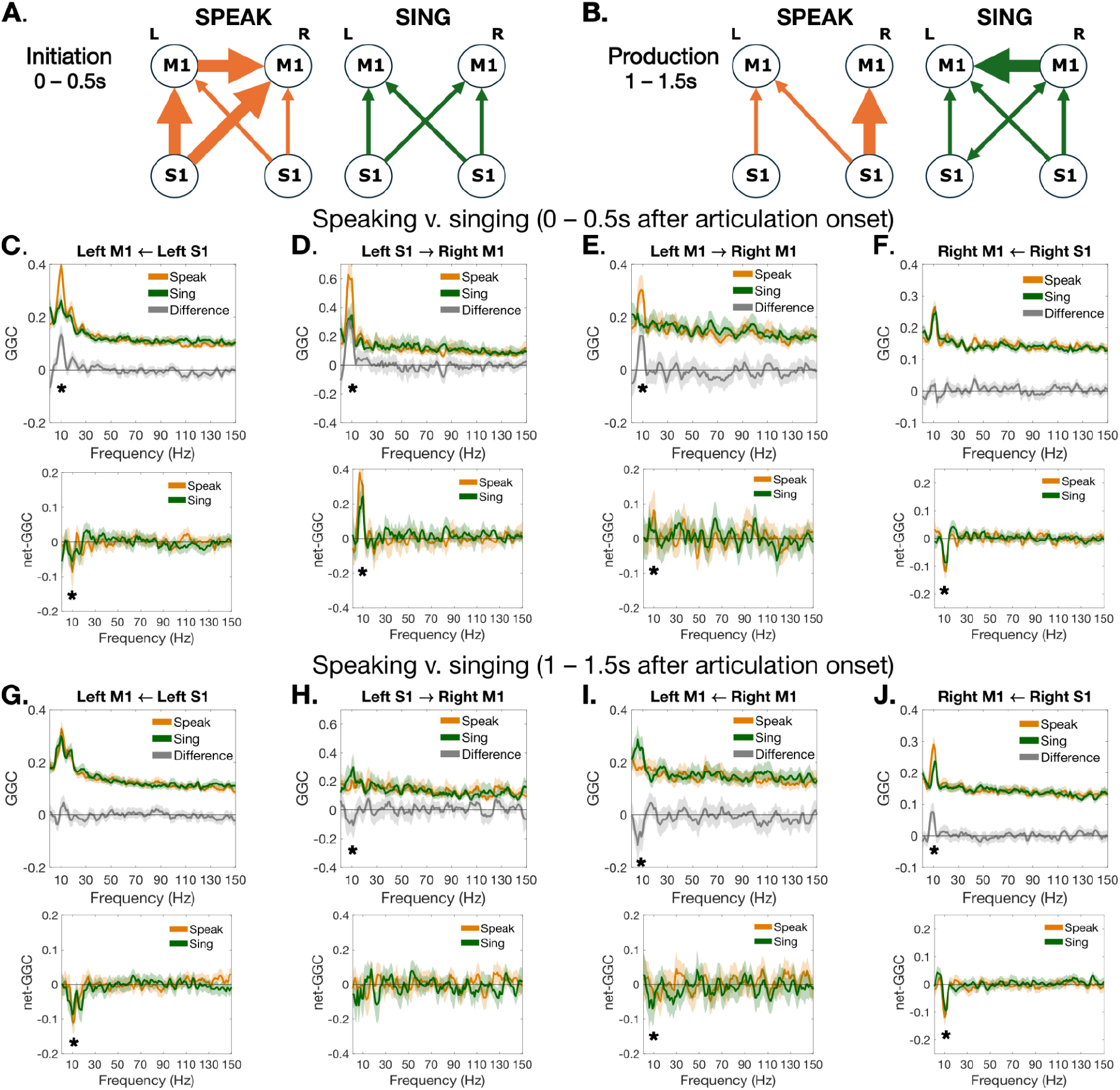
Granger causality analysis reveals a stronger left lateralization in the initiation of speaking. **A&B:** Schematic representation of the directed information flow between areas of the SMC network, in the speaking (orange) and singing (green) conditions, within 0.5 s post-articulation (A) and 1–1.5 s post-articulation (B). Thin lines represent significant net information flow from one area to another (as shown by each arrow), thick arrows represent significant information flow and difference in strength of information flow between condition (e.g., speaking > singing). **C–F:** Results of the nonparametric Granger– Geweke Causality (GGC) analysis in the ‘initiation’ interval (0–0.5 s after articulation onset; *top*) are shown together with the net influences in the represented inter-areal connection (*bottom*). *Top*: Frequency-domain nonparametric GGC for speaking (orange) and singing (green), together with the difference between the two (gray). Black asterisks represent a significant difference between conditions in the mu-band (8–12 Hz). Titles of each plot represent directionality with arrows from source to target (e.g. Left M1 → Right M1 plots functional connectivity from left M1 to right M1 whereas Left M1 ← Right M1 displays the opposite directionality). *Bottom*: Net influence for speaking (orange) and singing (green), obtained for each condition as the difference between the connectivity from area 1 → area 2 minus the connectivity from area 2 → area 1. Positive values indicate a stronger directed influence from area 1 → area 2 than vice versa; the opposite for negative values. Here, black asterisks indicate that the net influence in the mu-band (8–12 Hz) was significantly different from zero for at least one of the two conditions. In each panel, shadings represent 95% confidence intervals (CIs) across the electrode pairs for that connection. Only the dominant connections are plotted in this figure. **G–J:** Same as in C–F for the results obtained in the ‘production’ interval (1–1.5 s post-articulation).

Together, this pattern reveals that initiation in speaking is organized by left S1, which directly drives motor activity in the left M1 and right M1. Singing also engages this sensory-to-motor flow, but does not have an initial hemispheric bias, suggesting flexible initiation (see Fig. 5A. for schematic).

#### Production: 1–1.5 s post-articulation

In both speaking and singing, left S1 was leading left M1, (speaking: *t*(548)=-7.2215, *p*<0.0001); singing: *t*(548)=-6.1907, *p*<0.0001) (Fig. 5G), but there was no significant difference between conditions in net influence direction (left M1 → left S1: *t*(548)=1.0259, *p*=0.3054; left S1 → left M1: *t*(548)=1.8611, *p*=0.0633). Between the left S1 and right M1, connectivity was higher in singing (left S1 → right M1: *t*(69)=-3.4174, *p*=0.0011; right M1 → left S1: *t*(69)=-3.8833, *p*=0.0002), but net influence direction from left S1 → right M1 was not significantly different (speaking: *t*(69)=-0.8509, *p*=0.3978; singing: *t*(69)=-1.1915, *p*=0.2375) (Fig. 5H), which indicates increased bidirectional connections between the left S1 and right M1 in singing (Fig. 5B). Connectivity was higher for singing in the connections from right M1 → left M1 (*t*(189)=-2.7592, *p*=0.0064) (Fig. 5I). Connectivity from left M1 → right M1 was not significantly different between conditions (*t*(189)=-1.9165, *p*=0.0568). We found a stronger net influence direction from right M1 → left M1 in the singing condition (*t*(189)=-2.8962, *p*=0.0042), but not in speaking (*t*(189)=-1.6121, *p*=0.1086) (Fig. 5I). Connectivity was higher for speaking both in the connections from right S1 → right M1 (*t*(870)=4.9012, *p*<0.0001) and in the connections from right M1 → right S1 (*t*(870)=4.4158, *p*<0.0001) (Fig. 5J). We found a stronger net influence direction from right S1 → right M1 in both conditions (speaking: *t*(870)=-8.6830, *p*<0.0001; singing: *t*(870)=-8.2860, *p*<0.0001). Additionally, we observed net influence direction from right S1 → left M1 in both conditions (speaking: *t*(142)=-3.4849, *p*=0.0007; singing: *t*(142)=-2.3530, *p*=0.0200) (Fig. 5B). However, we did not see a significant difference in functional connectivity between conditions (left M1 → right S1: *t*(142)=-0.6327, *p*=0.5279; right S1 → left M1: *t*(142)=0.5375, *p*=0.5918).

As production unfolds, singing showed a distinctive interhemispheric motor net influence direction from right M1 → left M1, and a generally preserved right sensory-to-motor flow, consistent with a right-led, bilaterally-coordinated route that progressively recruits left SMC (see Fig. 5B. for schematic). Speaking showed an asymmetry that is distinct from its left-biased initiation, with net influence direction from right S1 → right M1 significantly stronger in speaking than singing later in production.

### An integrative mechanism for singing production

The current study provides evidence that singing and speaking engage a shared mu-coupled sensorimotor substrate through different network dynamics: a left-SMC mu rhythm that locally organizes HFA across both behaviors, intra- and interhemispheric mu-synchrony that ramps up in singing, and increased HFA in the right SMC for singing, together providing a route for production of linguistic content through song (Fig. 6). Our findings indicate that both speaking and singing require interhemispheric interaction but engage the network differently at initiation and throughout production. In speaking, early mu-band directionality originates in left S1, driving left M1 and right M1 and the interaction between the motor strips, accompanied by right S1 → right M1 sensory-to-motor flow. Speaking is left-led but bilaterally executed, a pattern that corroborates the long-standing clinical observation that left-hemisphere injury is typically necessary to produce aphasia, even though successful speech depends on interactions across both hemispheres.

**Figure 6.**
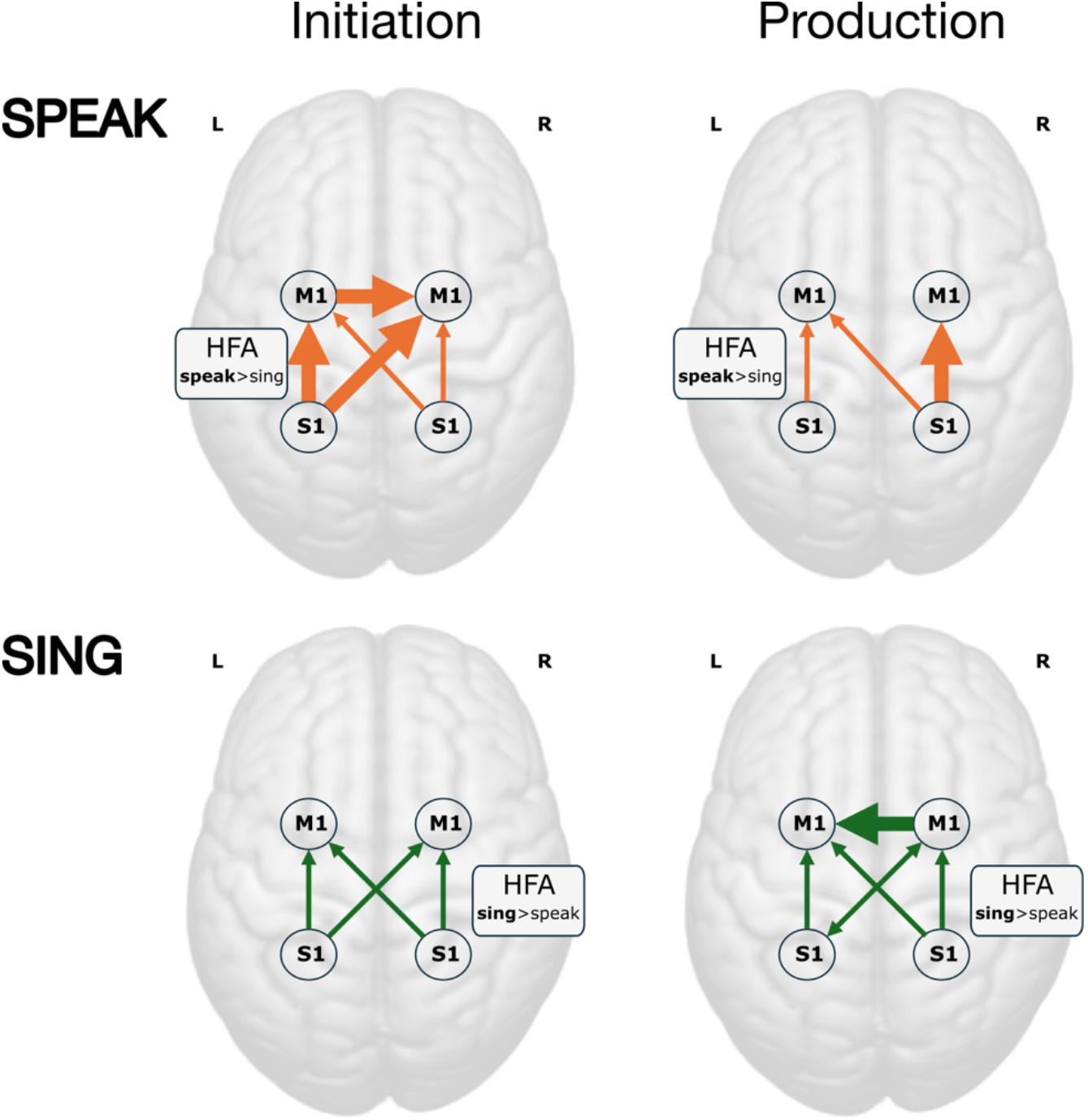
Schematic of control mechanism for initiation and production of singing vs. speaking. The schematic depicts the differential network recruitment of a shared SMC mu-coupled substrate during initiation and production of singing versus speaking. Initiation (0–0.5 s post-articulation onset) and production (1–1.5 s post-articulation onset) are shown in the *left* and *right* panels, respectively. Circles mark S1 and M1 in each hemisphere; arrows depict mu-band (8–12 Hz) synchrony directionality; bolded arrows represent stronger connectivity for a specific condition; inset boxes summarize HFA lateralization (speak > sing in the left SMC; sing > speak in the right SMC). **Top:** In speaking, initiation originates from left S1, which influences left and right M1 via mu-band modulation, and is accompanied by right S1 to right M1 flow, yielding a left-led but bilaterally engaged network. During production, mu-synchrony shifts directionality to the right hemisphere, with right S1 dominating influence to right M1, and no exchange between the motor strips. HFA is lateralized to the left SMC for both initiation and production. **Bottom:** In singing, initiation originates from both left S1 and right S1, but without a strong lateralization of influence. During production, right M1 leads left M1 with causal influence in the mu-band. HFA is lateralized to the right SMC for both initiation and production. Singing engages a more flexible, less left-lateralized, right-assisted network that may be more adaptable to motor output after left-hemisphere injury.

By contrast, singing does not require a stronger left-hemisphere initiation at onset. Early directionality lacks dominance from the left or right motor strip and is marked by right S1 → right M1 drive (present in both speaking and singing). Right-lateralized HFA supports local execution, and a net influence from right M1 → left M1 emerges as production unfolds. In parallel, singing shows a ramp-up of mu-synchrony within left S1–M1 and between the two M1s, indicating progressive recruitment of left-hemisphere circuitry via interhemispheric coupling. The same left-SMC mu–HFA coupling architecture is present in both behaviors, suggesting that what differs between singing and speaking is not the existence of this local substrate but how it is recruited across the network as a sentence unfolds.

For vocal production, the end goal in both singing and speaking is motor execution in M1. Speaking reaches this goal predominantly via left S1 feeding left M1 (left-biased HFA, early left-led mu-directionality). Singing can reach M1 through a right-led route, with right S1 influencing right M1, right-hemisphere HFA indexing execution, and causal influence from right M1 to left M1 that brings the left SMC online (see Fig. 6 for a schematic of this integrative mechanism).

Taken together, singing provides a more flexible pathway: it can be initiated through left or right-hemisphere sensory-motor channels and then enlists the left SMC through strengthened interhemispheric interactions, rather than relying on a strictly left-initiated trigger. This alternate engagement of the left and right SMC is a possible route for preserved language to be expressed, even in cases of left hemisphere injury. The preservation of interhemispheric communication routes, and right hemisphere integrity is critical. For instance, not all individuals with aphasia from left hemisphere injury are able to express themselves fluently in song^41,42^. Aphasia results from lesions that damage the middle cerebral territory, compromising the white matter connections within and between the two hemispheres^43–45^. Of note, nearly all individuals who stutter have lesion-free hemispheres, and show the ability to sing without a stutter^2,3^.

## Discussion

### A neural mechanism for the expression of linguistic content through song

Intracranial EEG recordings revealed that HFA power is right-lateralized for singing sentences in the sensorimotor cortex and left lateralized for speaking. We also found distinct dynamics for singing versus speaking within the mu-band (∼10 Hz), a prominent rhythm observed in the sensorimotor cortex in both hemispheres. Phase-amplitude coupling analyses further revealed that mu phase locally organizes HFA within left SMC in a channel-specific, spatially interdigitated coupling architecture that is preserved across both behaviors, indicating that speaking and singing share a common sensorimotor substrate that is differentially recruited across the network. During singing, high-frequency activity (HFA, 70–150 Hz) was right-lateralized, and mu-band (∼10 Hz) synchrony ramped up over time within left SMC and between the two M1s, indicating progressive recruitment of left circuitry via interhemispheric cooperation. In speaking, HFA was left-lateralized, and mu-band dynamics were left-led at onset with sustained right-hemisphere involvement thereafter. Directional analyses further showed that speech initiation reflects left S1 → left/right M1 drive, whereas singing does not require left-hemisphere initiation at onset and instead can proceed through right S1 → right M1 or left S1 → right M1 drive followed by right M1 → left M1 influence as production unfolds. Together, these findings support a model in which singing and speaking engage a shared mu-coupled sensorimotor substrate through distinct recruitment dynamics: speaking via a left-led, bilaterally executed route, and singing via a right-hemisphere led, bilaterally-coordinated route that progressively recruits left sensorimotor nodes via interhemispheric synchrony.

In our data, we observe a ∼10 Hz rhythm with a ∼20 Hz harmonic, classically described as a mu rhythm, that coordinates left–right interactions in S1/M1. Work by Norman et al. (2025) suggests that a closely related 6–10 Hz “sensorimotor theta” rhythm is an intrinsic organizing frequency of the speech motor system: during articulation, bilateral SMC shows enhanced phase coherence in this band, coupled to bursts of high-frequency activity and to pulse-like vocal-tract kinematics at specific phases^16^. Our phase-amplitude coupling results provide direct, channel-resolved evidence that concurs with the multiplexing proposed by Norman and colleagues: mu-band phase locally organizes HFA in left SMC, with channel-specific phase preferences that are stable across trials, spatially interdigitated, and preserved across speaking and singing. This observation, together with prior ECoG work showing functionally specific but spatially intermixed representations in sensorimotor speech cortex^10,12^, supports the idea that speech production depends on precise phase alignment of an intrinsic SMC oscillation that multiplexes sensory feedback, motor commands, and articulator dynamics. In Broca’s aphasia, where lesions typically involve left ventral peri-rolandic and insular cortex and their underlying white matter^44^, this intrinsic generator and its coupling pathways are disrupted. The result is a system in which local rhythms can be expressed, but their phases fail to align reliably across the network, consistent with halting, effortful speech, variable apraxic errors, and occasional “lucky” error-free productions of complex words.

Within this framework, singing provides an alternative way to engage the same intrinsic rhythm in a more stable manner. The rhythmic–melodic scaffold imposes an external temporal structure that can reinforce mu/theta-band synchrony within and between hemispheres and tighten the alignment between oscillatory phase, HFA bursts, and vocal-tract kinematics. Our results show that singing preferentially enhances right-hemisphere HFA and mu-band coupling and promotes a right-to-left motor influence as sentences unfold. This pattern offers a mechanistic account of the classic clinical dissociation in Broca’s aphasia: even when left-hemisphere regions that normally initiate speech are damaged, spared peri-rolandic tissue and intact right SMC can still participate in a right-led, bilaterally-coordinated oscillatory regime that supports more reliable sequencing of articulatory gestures in song than in speech.

### Feedforward and feedback control of speech and song

The left and right sensorimotor cortices jointly subserve planning, execution, and monitoring of articulatory movements. In the Directions into Velocities of Articulators model (DIVA)^39^ framework, feedforward speech initiation commands are primarily routed through the left motor cortex, with the right motor cortex contributing more to feedback control after movement onset. Consistent with this model, and also with a recent MEG report^46^ that lateralization shifts across time in the mu rhythm from left to right in speaking, our intracranial data also show early left-biased mu-synchrony during speaking that evolves into sustained right-hemisphere involvement, alongside left-lateralized HFA. By contrast, singing does not exhibit a left-initiated onset; instead, mu-band coupling ramps within left S1–M1 over time while right-lateralized HFA and right-to-left influences support execution, suggesting a different feedforward/feedback balance for song.

### A shared sensorimotor substrate, differentially recruited

A central implication of our findings is a distinction between the architecture of sensorimotor cortex and the way it is engaged. The phase-amplitude coupling results demonstrate that mu phase locally organizes HFA in left SMC in a channel-specific, spatially interdigitated fashion, and that this local coupling is present in both speaking and singing. The mu-synchrony and directionality results, by contrast, reveal that singing and speaking differentially recruit this substrate at the network level, through distinct initiation biases, interhemispheric coupling dynamics, and directional information flow. This architecture-versus-use distinction reframes the clinical dissociation between singing and speaking: rather than positing entirely separate circuits, it suggests that song and speech share a common sensorimotor substrate whose recruitment can be flexibly modulated by the structural scaffolding of music.

### Implications for future brain-computer interfaces (BCI)

Recent brain-computer interfaces have successfully used HFA or spiking obtained from dense electrocorticographic grids or Utah arrays in the left primary motor cortex to decode speech in individuals with paralysis or anarthria^11,47,48^. These approaches in left M1 capture phoneme-specific content that, when combined with predictive language models, can be used to decode speech from an intact cortex despite paralysis of the vocal-motor system. Our results showed that HFA tracks local execution of motor commands in the left SMC for speaking with an increase in HFA in the right SMC for singing, supporting enhanced population activity. In addition to the HFA findings, we observed differences in local and interhemispheric synchrony in singing versus speaking. We find that mu-band activity indexes intra- and interhemispheric communication between left and right M1 and S1 with differential pre- and post-articulatory dynamics as the sentence unfolds. Bilateral SMC synchrony increases at the onset of speaking, while this synchrony continues to ramp up in singing. In addition, our phase-amplitude coupling analysis revealed that individual left-SMC contacts carry stable, discrete mu-phase preferences that are consistent across trials and across speaking and singing. This channel-specific phase tuning suggests that not only mu amplitude, but also mu phase, could serve as an additional feature space for decoding, offering a low-dimensional code that may be robust across production modes. Rapid sentence-level production is the ultimate goal of a speech prosthetic. Future BCI models could use mu-band activity and phase-tuning in both the left and right sensorimotor cortex as a state-signal to modulate decoder weights as a sentence progresses and enhanced singing could be obtained by selective mu band neuromodulation. Bilateral invasive or non-invasive electrodes over SMC could be used to boost singing or speaking capacity in patients with disabling speech output problems.

An important challenge for the field is to create a BCI for individuals with language impairments from aphasia or apraxia of speech, a motor speech disorder that affects the ability to coordinate complex articulatory sequences. A primary difficulty is selecting the neural substrate for decoding or modulating in individuals with an injured cortex. Behaviorally, singing allows expression through alternate routes in an impaired speech and language system^2,4,5^. Here, we show that the neural mechanisms of speech and song engage the left and right sensorimotor cortices in distinct ways within the mu-band and in HFA power modulations. Incorporating these findings from singing into training a BCI could be employed to develop a speech prosthetic for individuals with a compromised speech and language production system.

## Methods

### Patients

We collected intracranial EEG data from 18 patients (mean age 35.2 +/-14.3 years; 11 females) undergoing intraoperative neurosurgical treatment for pharmacoresistant epilepsy. All patients were implanted with stereo EEG depth electrodes to determine the site of seizure onset for surgical resection. We selected 13 patients, without lesions, that had coverage in the left or right sensorimotor cortex. The electrode placement was managed entirely by the neurosurgical team for each patient. The recordings took place at multiple hospitals including Loma Linda University Medical Center (n = 1), Barnes-Jewish Hospital in St. Louis (n = 11), and the St. Louis Children’s Hospital (n = 1). All patients provided written informed consent as part of the research protocol approved by each hospital’s Institutional Review Board and by the University of California, Berkeley. Patients were tested when they were fully alert and willing to participate.

Our cohort involved individuals with epilepsy undergoing presurgical monitoring, which may affect the generalizability of the findings. We minimized disease- and treatment-related confounds by testing only when patients were fully alert and by excluding contacts with epileptiform activity or other artifacts. Electrode implantation was dictated by clinical need rather than experimental design, yielding uneven, asymmetric coverage across participants; consequently, some effects may have gone undetected due to limited sampling.

### Behavioral task

Participants were presented visually and auditorily with matched sentences and were asked to repeat them once as quickly as possible. The stimuli were adapted from a standard motor speech assessment battery^49^, designed to engage motor speech planning and execution skills. The stimuli contained 5-7 words, some of which were multisyllabic with consonant clusters that required shifting between points of articulation. For example, in the word ‘spaghetti’, there is an initial consonant cluster, which demands quick movement between distant points of articulation (bilabial /p/ to velar /g/ and back to apico-alveolar /t/). An example sentence from the stimuli used is “prepping lobster is the best”. A block design was used for singing and speaking conditions and the order of presentation was counterbalanced across participants. To optimize for a hospital setting, in which time is limited and patients tire quickly, 20 sentences were presented in each of the spoken and sung conditions, for a total of 40 trials.

### Data acquisition

Electrophysiological data were collected using BCI2000, an open-source software, at each hospital in the epilepsy monitory unit^50^. The sampling rate was 512 Hz at Loma Linda Hospital, and 2000 Hz at Barnes-Jewish Hospital and Saint Louis Children’s Hospital.

### Anatomical reconstructions

An anatomical data processing pipeline was used to localize electrodes from a pre-implantation MRI and a post-implantation CT scan^51^. The MRI and CT images were co-registered using a rigid body transformation and adjusted for brain shifts during implantation and surgery. The FreeSurfer analysis suite^52^ was used to generate a native-space reconstruction of the patient’s brain with electrodes visualized in precise anatomical locations. Electrodes were classified by a board-certified neurologist according to the anatomy in each patient’s native space. Only electrodes identified in the regions of interest for the study were included in the analysis: IFG (left: 34 electrodes, right: 41 electrodes), STG (left: 42 electrodes, right: 21 electrodes), PostCG/S1 (left: 41 electrodes, right: 47 electrodes), PreCG/M1 (left: 44 electrodes, right: 47 electrodes), MFG (left: 30 electrodes, right: 62 electrodes), SMG (left: 5 electrodes, right: 10 electrodes). For visualization of coverage across participants, we converted native-space electrodes reconstructions to Montreal Neurological Institute (MNI) space, using volume-based normalization. Fig. 1A shows the electrodes coverage across participants.

### Data preprocessing

Continuous iEEG data were low-pass filtered at 150 Hz and notch-filtered at 60 Hz and harmonics (up to 120 Hz) to remove line noise. Signals were re-referenced using a bipolar montage. Channels with poor contact or excessive noise were excluded after visual inspection, and a neurologist identified and removed electrodes exhibiting epileptiform activity or epochs contaminated by spread from the seizure focus. Data were epoched relative to the event of interest (articulation, see below). All signals were downsampled to 400 Hz to account for variable sampling rates across sites. All preprocessing was conducted in Python using the MNE toolbox package^53^. Custom MATLAB codes were used for analysis of the preprocessed data.

### Manual articulation tagging and alignment

Audio was recorded synchronously with iEEG for each trial for all patients. For every waveform, trained raters manually annotated speech onset in Praat^54^ from the waveform and spectrogram, creating item-level transcriptions time-aligned to the onset of articulation. Event markers from the acquisition/BCI system were then used to synchronize audio and neural streams and to verify timing consistency. The resulting onset timestamps were exported and used to generate articulation-locked epochs for all analyses.

### Irregular-Resampling Auto-Spectral Analysis (IRASA)

To estimate the fractal component of the power spectrum, we applied IRASA^29^ to the broadband signal by resampling the time series with a set of non-integer factors h (typically spanning 1.1– 1.9), using both upsampling (h) and the corresponding downsampling (1/h). For each factor, we computed the auto-power spectra of the two resampled signals and combined them via their geometric mean to obtain a factor-specific estimate of the fractal spectrum. Because non-integer resampling shifts narrowband oscillatory power to h-dependent frequencies while leaving the fractal component largely invariant, we aggregated these factor-specific fractal estimates by taking the median across h at each frequency, yielding a robust estimate of the fractal PSD. The oscillatory PSD was then derived by subtracting the fractal PSD from the mixed PSD; to reduce residual cross-terms, PSDs were additionally averaged across multiple time segments assumed to be stationary, under the assumption of no systematic phase coupling between fractal and oscillatory components.

### Time–frequency power estimates and cluster-based permutation (ROI-level)

For each participant and condition (speak, sing), iEEG was epoched articulation-locked and from −3 to +2 seconds. Time–frequency power was computed with a Morlet wavelet transform (custom MATLAB; FieldTrip^55^) at frequencies from 1–150 Hz in 1-Hz steps, using *ω*_*0*_=6 and zero-padding. The resulting power estimates were grouped for the different ROIs using the information about electrodes labels pulled from the anatomical descriptions (see Anatomical reconstructions for details, e.g., Left/Right PreCG, PostCG). Prior to statistics, time-varying power estimates were smoothed with a 41-sample moving average along the time axis.

Condition differences (speak vs. sing) were tested within each ROI using FieldTrip’s cluster-based permutation approach, using a two-tailed dependent-samples *t*-statistic (with *p*<0.05, alpha level distributed over both tails by multiplying the *p*-values with a factor of 2 prior to thresholding), 10,000 randomizations, and *p*_*perm*_<0.05 for the permutation test. The comparison was restricted to -0.5–2 s interval around articulation onset for inference. Tests were performed over time– frequency bins (no channel-neighbor constraint), and significant effects are reported as cluster-corrected regions.

### Time-varying intersite phase clustering (ISPC)

For each participant and condition (speak, sing), iEEG trials were articulation-locked (task-specific windows); when needed, trial counts were equalized within-subject by random subsampling. Contacts were anatomically labeled and analyses were restricted to pre-specified ROIs (PreCG/M1, PostCG/S1; left/right). Time–frequency decomposition used a Morlet wavelet (custom MATLAB; FieldTrip^55^) using the same parameters adopted for power estimation (frequencies 1–150 Hz in 1-Hz steps, *ω*_*0*_=6, zero-padding). From the complex coefficients, phase angles were extracted and intersite phase clustering (ISPC)^34^ was computed for every channel pair, time point, trial, and frequency, then averaged across trials to yield time-resolved pairwise synchrony. The number of electrode pairs on interareal connections of interest were as follows: left M1–left S1 (n=549), left M1–right S1 (n=143), right M1–right S1 (n=871), left M1–right M1 (n=190), left S1–right M1 (n=70). None of the participants had simultaneous coverage in left S1– right S1.

### Cluster-based permutation on ISPC (ROI↔ROI)

Group inference targeted predefined ROI-to-ROI links (e.g., left M1–left S1; left M1–right M1). For each participant, we stacked all electrode pairs matching a given connection and extracted their ISPC within 1–70 Hz and −1.5 to +2 s around articulation; statistics were performed over the interval -0.5–2 s. A 41-sample moving average was applied before testing. Sing vs. speak contrasts used FieldTrip’s cluster-based permutation approach, using a two-tailed dependent-samples *t*-statistic (with *p*<0.05, alpha level distributed over both tails by multiplying the *p*-values with a factor of 2 prior to thresholding), 10,000 randomizations, *p*_*perm*_<0.05 for the permutation test, no channel-neighbor constraint, and electrode pairs as the repeated unit in the design. Significant time–frequency clusters are reported as cluster-corrected effects. Mu-band (8–12 Hz) ISPC was averaged over frequency and converted to relative synchrony change (RSC) by z-scoring against a -0.5–0 s pre-articulation baseline within each condition, to display condition-wise time courses with confidence intervals.

### Granger causality analysis

Granger causality provides an operational definition of directed functional influence between two time series^56^. In this framework, activity in region X is said to causally influence activity in region Y if incorporating the past of X improves prediction of the current (or future) values of Y beyond what can be achieved from Y’s own past (and, when modeled, the past of other observed signals). Under a linear, stochastic dynamical model, this predictive improvement corresponds to a reduction in the residual variance (prediction error) of Y when X is included as a predictor, and can be expressed as an asymmetric measure of directed dependence.

In the frequency domain, Granger–Geweke causality (GGC) decomposes this directed predictability by frequency, quantifying how strongly fluctuations in X at each frequency contribute to predicting fluctuations in Y at the same frequency^40^. Practically, the method is implemented by estimating the cross-spectral density of the multichannel data and applying spectral factorization to obtain the transfer function and noise covariance that define the underlying linear system. This formulation is well suited for oscillatory neural signals because it allows directionality to be evaluated within targeted frequency bands (here, the mu band), and it distinguishes directional influence (X → Y) from the reverse direction (Y → X) within the same data segment, conditional on the signals included in the model.

We performed a connectivity analysis using the Granger causality framework, to test for directionality of the influences between areas of the SMC network, in two time intervals (‘initiation’: 0–0.5s; ‘production’: 1–1.5s post-articulation). In each interval, we derived a frequency domain measure of GGC between two areas^39^. For this, we used a multitaper-based approach for cross-spectral density estimation (time-bandwidth product *NW*=2) and nonparametric spectral factorization^57–59^. For each inter-areal connection, we compared the directed connectivity between speak and sing in the mu-band (8–12 Hz) using a paired-samples *t*-test across all electrode pairs for that connection (*df* = number of electrode pairs - 1). In the same connection, the dominant directionality of the influences was tested by using a *t*-test on the net influence (area 1 → area 2 minus area 2 → area 1) across all electrode pairs (*df* = number of electrode pairs - 1). If the net influence was significantly higher than zero it indicates that the directed influence from area 1 → area 2 is stronger than the directed influence from area 2 → area 1; vice versa, when the net influence was significantly lower than zero.

### Phase-amplitude coupling (PAC)

Phase-amplitude coupling (PAC) was quantified using Tort’s modulation index (MI)^33^ within the articulation-locked production window (0 to +2000 ms after articulation onset), separately per electrode and condition (sing, speak). Within each participant, sing and speak trials were randomly subsampled to the smaller condition’s count to equate trial numbers prior to MI estimation. The MI was estimated across a frequency-frequency grid spanning phase frequencies of 2–20 Hz (1 Hz steps, 2 Hz bandwidth) and amplitude frequencies of 30–150 Hz (5 Hz steps, bandwidth equal to twice the phase center frequency), using 18 phase bins. Epochs were extended by 500 ms on each side of the analysis window and trimmed after filtering to prevent edge artifacts. To standardize MI values and assess significance, following the approach of Daume et al.^60^, 200 surrogate MI distributions were generated per channel by randomly shuffling the phase-amplitude pairing across trials. A normal distribution was fit to each surrogate distribution and observed MI values were z-scored accordingly (MI(z)). A channel was considered to show significant mu–HFA PAC when its mean MI(z) within the band of interest (8–10 Hz phase × 70–150 Hz amplitude) exceeded z = 1.64 (one-tailed, p < 0.05), averaged across conditions; the proportion of channels with significant PAC was assessed per ROI × hemisphere. In a separate analysis, we further derived single-trial MI estimates (no surrogates) and binned single-trial HFA amplitude (70–150 Hz) into the 18 mu-phase bins (8–10 Hz) to visualize trial-by-trial PAC dynamics and characterize mu-phase preferences in individual participants and channels.

## Data Availability

The data that support the findings of this study are available on request from the corresponding author [ALP]. The data are not publicly available due to patient privacy.

## Acknowledgements

This work was supported by the National Institutes of Health (Grants R21DC021042 and R01DC016345 to NFD; Grants R01NS021135 and P50MH109429 to RTK).

## Notes

### Competing Interest Statement

The authors have declared no competing interest.

### Summary of Updates

Added phase-amplitude coupling analysis; figure 3 revised.

